# Spillover effects of a combined water, sanitation, and handwashing intervention in rural Bangladesh: a randomized controlled trial

**DOI:** 10.1101/191726

**Authors:** Jade Benjamin-Chung, Nuhu Amin, Ayse Ercumen, Benjamin F Arnold, Alan Hubbard, Leanne Unicomb, Mahbubur Rahman, Stephen P Luby, John M. Colford

## Abstract

**Background:** Water, sanitation, and handwashing (WSH) interventions may confer indirect benefits (“spillovers”) on neighbors of recipients by interrupting pathogen transmission. We measured geographically local spillovers in WASH Benefits, a cluster-randomized trial in rural Bangladesh, by comparing outcomes among neighbors of intervention vs. control participants.

**Methods:** WASH Benefits had randomly allocated geographically-defined clusters to a compound-level intervention (chlorinated drinking water, upgraded sanitation, and handwashing promotion) or control and followed children for two years. We enrolled neighboring children age-matched to trial participants that would have been eligible for WASH Benefits had they been conceived slightly earlier or later. After 28 months of intervention, we quantified fecal indicator bacteria in toy rinse and drinking water samples, measured soil-transmitted helminth infections, and recorded caregiver-reported diarrhea and respiratory illness. Neither fieldworkers nor participants were masked. Analysis was intention-to-treat.

**Results:** We enrolled neighbors of WASH Benefits participants in 90 control (N=900) and 90 intervention clusters (N=899). Neighbors’ characteristics were balanced across arms. The prevalence of any detectable *E. coli* in tubewell samples was lower for neighbors of intervention vs. control (prevalence ratio=0.83; 0.73, 0.95). There was no difference in *E. coli* and coliform prevalence between arms for other environmental samples. Disease prevalence was similar in neighbors of intervention vs. control participants: *Ascaris* (prevalence difference [PD]=0.00; -0.07, 0.08), hookworm (PD=0.01; -0.01, 0.04), *Trichuris* (PD=0.02; -0.02, 0.05), diarrhea (PD=0.00; -0.02,0.03), respiratory illness (PD=-0.01; -0.04, 0.03).

**Conclusions:** We found spillover effects of a compound-level combined WSH intervention for tubewell water contamination but not for child health outcomes.

**Key Messages:**

- Water, sanitation, and handwashing (WSH) interventions may confer indirect benefits (“spillovers”) on neighbors of recipients by interrupting pathogen transmission, reducing environmental contamination, or spurring the adoption of health behaviors.
- We conducted a randomized trial in rural Bangladesh to measure whether neighbors of a compound-level WSH intervention improved hygiene behaviors and had lower prevalence environmental contamination, soil-transmitted helminth infection, diarrhea, and respiratory illness among children under 5 years after two years of intervention.
- We did not find evidence of intervention adoption or improved hygiene behavior among neighbors of a WSH intervention delivered for 2 years.
- The WSH intervention reduced fecal contamination of neighbors’ tubewell water but did not lead to spillovers for other proximal measures of contamination in the domestic environment or for child health outcomes. For proximal spillover effects to translate to distal spillover effects, improvements in neighbors’ health behaviors may have been necessary.

## Introduction

Improvements in household water quality, handwashing practices, and sanitation (WSH) may reduce the risk diarrhea,^1^ soil-transmitted helminth (STH) infection^2^ and respiratory illness.^3^,^4^ WSH interventions may also reduce illness among neighbors through “spillover effects”^5^ (a.k.a. “herd effects”^6^–^8^ or “indirect effects”^9^) through 1) reduced fecal contamination in the environment surrounding intervention recipients, 2) reduced pathogen transmission from intervention recipients to neighbors, or 3) adoption of promoted health behaviors by neighbors. If WSH interventions reduce illness among both recipients and other individuals, intervention effect estimates that ignore spillover effects will be underestimated.

There is a rich literature on herd effects of vaccines.^5^,^7^ The literature on spillover effects for other infectious disease interventions, such as school-based deworming^10^ and insecticide treated nets,^11^ is growing.^5^ While many empirical studies have measured WSH interventions’ effects directly on recipients,^1^–^3^ few have measured spillover effects of WSH;^12^–^19^ these studies used observational designs to measure spillovers, so spillover estimates may be susceptible to bias if there are systematic differences between individuals in close proximity to intervention and individuals serving as controls.

We measured spillover effects of a compound-level combined WSH intervention in an existing, large, rigorously designed trial: the WASH Benefits Bangladesh trial.^20^ This study measured whether compounds neighboring WASH Benefits intervention recipient homes had lower environmental contamination and their children had lower prevalence of STH, diarrhea, and respiratory illness compared to children neighboring controls.

## Methods

### Randomization

We performed a cluster-randomized trial building upon the WASH Benefits Bangladesh study,^21^,^20^ which was conducted in Gazipur, Mymensingh, Tangail and Kishoreganj districts of central Bangladesh. These areas were selected because they had low groundwater arsenic and iron (to avoid interference with chlorine water treatment) and no other WSH or nutrition programs. WASH Benefits randomly assigned clusters to: 1) drinking water treatment and safe storage, 2) sanitation, 3) handwashing, 4) combined water + sanitation + handwashing (WSH) 5) nutrition, 6) combined nutrition + WSH, and 7) control (no intervention).^21^ WASH Benefits investigators randomized treatment within geographic blocks containing adjacent clusters. An investigator at UC Berkeley (BFA) used a random number generator to randomly assign treatment or control within groups of geographically contiguous clusters. Clusters were separated by at least 1 kilometer to reduce the risk of between-cluster spillovers resulting from reductions in disease transmission or the adoption of interventions in the control group. The study found no evidence of spillovers from the intervention to the control group.^20^

We measured geographically local spillovers among neighbors of trial participants in 90 control clusters and 90 combined WSH intervention clusters in WASH Benefits. To measure spillovers, this study focused on the combined WSH intervention because we hypothesized that of all intervention packages in the trial it was most likely to have spillover effects (Figure 1). We selected control clusters where the main trial planned to collect environmental samples to coordinate data collection efforts and maximize comparability with the main trial. Because interventions included visible hardware, neither the outcome measurement team nor study subjects were masked to intervention assignment.

**Figure 1:**
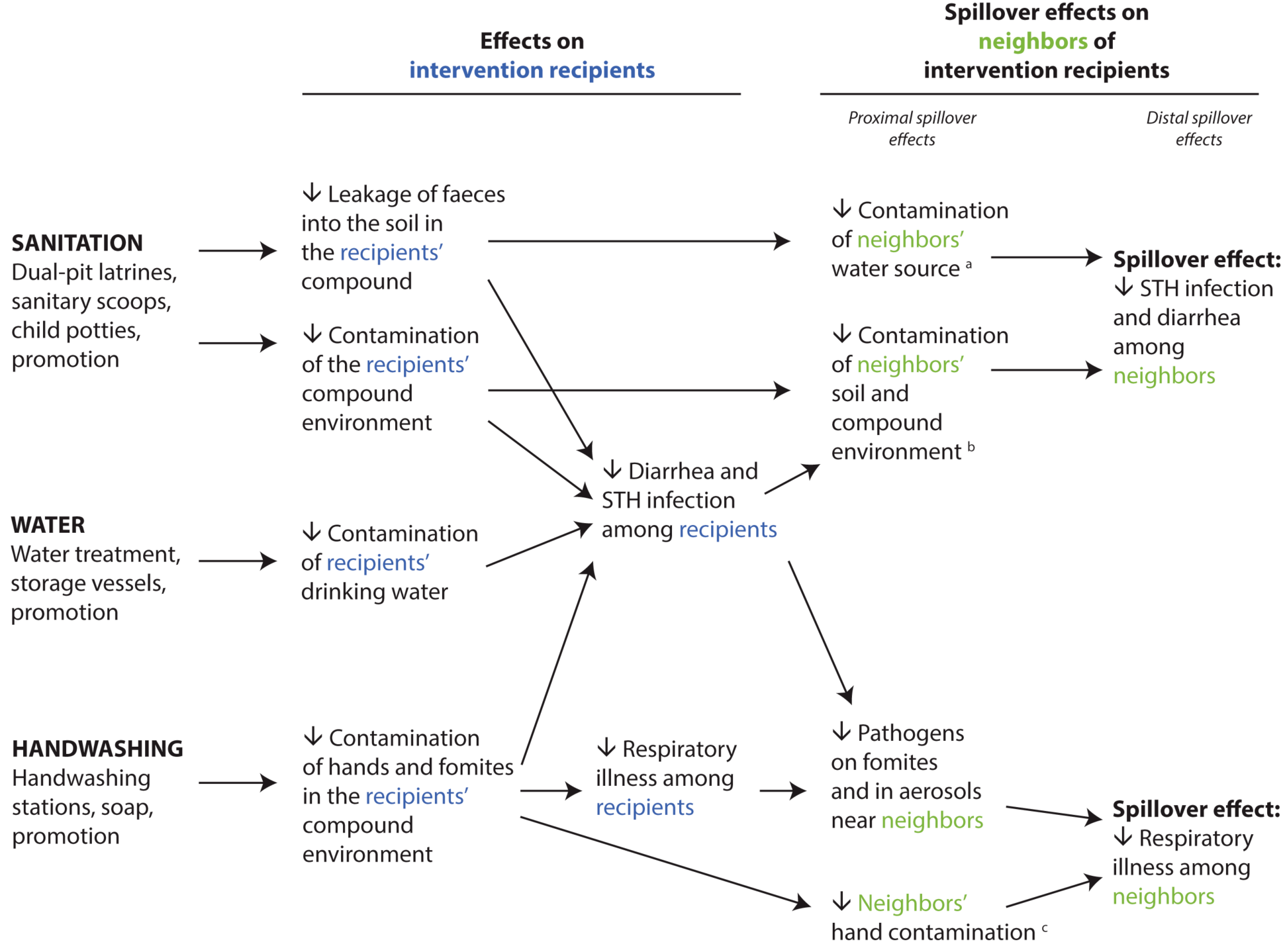
Theoretical model for spillover effects of a compound-level combined water, sanitation, and handwashing intervention. ^a^Contamination of drinking water with fecal indicator bacteria ^b^Synanthropic fly counts ^c^Observed hand cleanliness

### Participants

In rural Bangladesh, families typically live in clusters of households with a common courtyard. Compounds were eligible for WASH Benefits if a pregnant woman resided there at the time of study enrollment who intended to stay in their village during the follow-up period. The trial followed a birth cohort of “index” children *(in utero* at enrollment) of enrolled mothers. After 24 months of intervention, there were 6.4 study compounds per cluster on average, and these compounds typically comprised <10% of compounds located within the cluster boundaries. To measure spillovers, we enrolled neighboring compounds concurrent with primary outcome measurement in the main trial. Neighbors were eligible if a child 0-59 months at the time of spillover study enrollment (just younger or older than the index child cohort) resided there and if they were within 120 steps (2 minutes walking time) of a WASH Benefits compound (Figure 2). We excluded children enrolled in WASH Benefits and children in compounds that shared a courtyard, latrine, or handwashing station with a WASH Benefits compound. Within each cluster, we enrolled 10 compounds in the spillover study.

**Figure 2:**
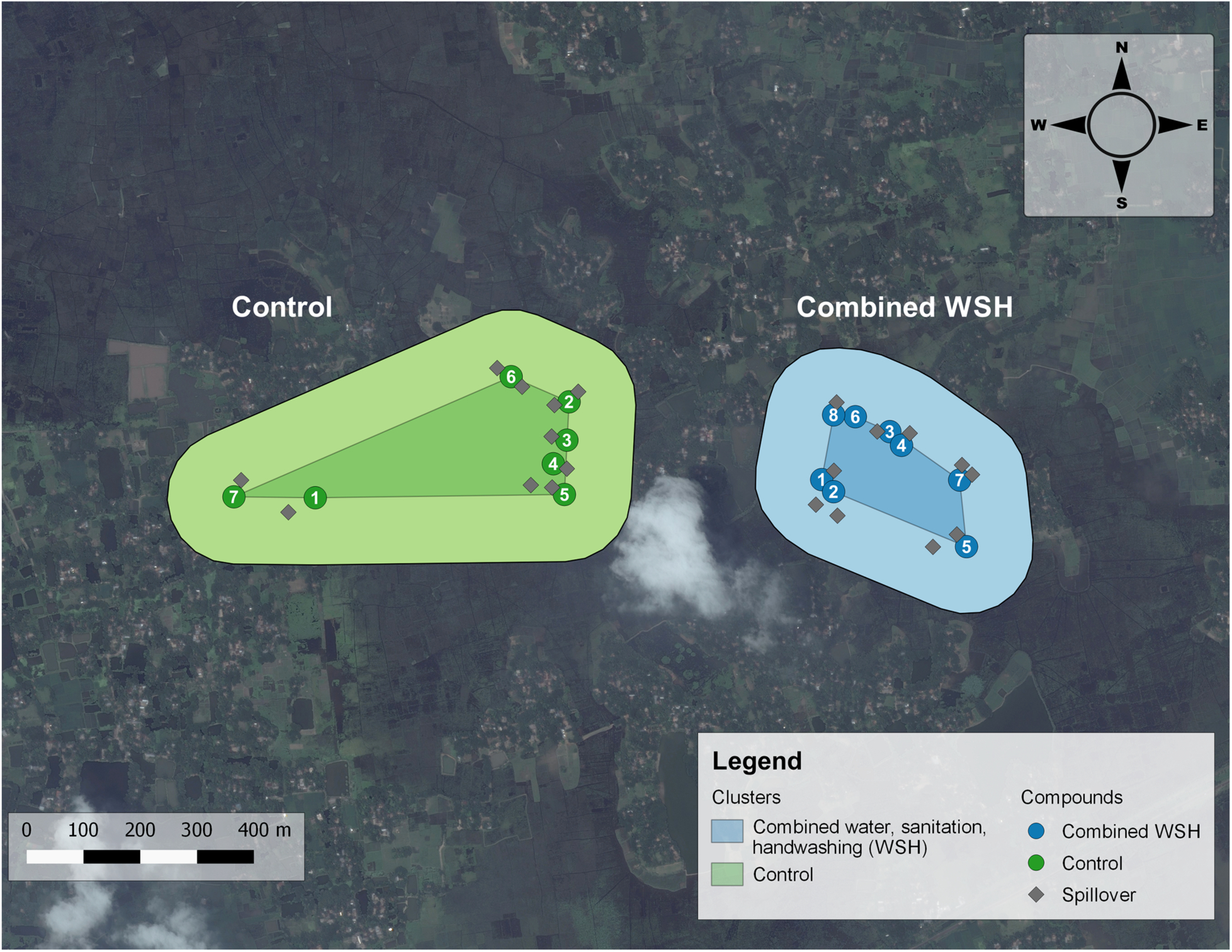
Study design. This figure depicts the study design in two clusters, one assigned to the combined water, sanitation, and handwashing intervention and the other assigned to control. Each cluster was separated by a buffer zone of at least 1 km to minimize the chance of spillovers between clusters. The numbered circles denote the compounds enrolled in the WASH Benefits study. The gray diamonds denote the neighboring compounds enrolled in the spillover study. The WASH Benefits study did not formally define the boundaries of each cluster. In this figure, the darker shaded center of each cluster is the polygon formed by linking the outermost compounds in each cluster, and the lighter shaded section is the periphery around this polygon. We restricted enrollment to the compounds within this periphery to ensure that the 1km buffer zone was maintained in this study.

### Interventions

Intervention recipients in the combined WSH arm received free chlorine tablets (Aquatabs®; NaDCC), a safe storage vessel to treat and store drinking water, child potties, sani-scoop hoes to remove feces from household, latrine upgrades to a double pit pour-flush latrine for all households in the compound; and handwashing stations including soapy water bottles and detergent soap. Local promoters visited study compounds on average once per week to encourage intervention uptake. The control arm and spillover study participants did not receive interventions or health promotion.

### Procedures

Fieldworkers administered a survey to caregivers of enrolled children at the time of enrollment into the spillover study, concurrent with primary outcome measurements in WASH Benefits (after 28 months of intervention). The survey measured household characteristics, child illness, WSH indicators (e.g., water treatment), and neighbors’ knowledge of the WASH Benefits and interactions with WASH Benefits participants and promoters.

Due to political instability in Bangladesh, environmental and biological samples for the spillover study and the WASH Benefits trial were collected 4 months after the survey (after 32 months of intervention) to ensure safe transport and a cold chain. If the child originally enrolled to measure spillovers was not present to provide a stool sample, we enrolled another child in the compound aged 0-5 years. All participants were offered a single dose of albendazole following stool collection regardless of infection status. Two slides were prepared from each stool sample and analyzed using Kato-Katz within 30 minutes of slide preparation.^22^ Laboratory technicians quantified *Ascaris*, hookworm, *Trichuris* ova on each slide. Counts were double-entered into a database by independent technicians. 10% of slides were counted by two technicians, and 5% were counted by a senior parasitologist for quality assurance.

In a subset of 86 control and 80 intervention clusters, fieldworkers collected drinking water samples and recorded water source (tubewell, stored water container or filter, tap). They distributed a non-porous, sterilized toy ball to each enrolled child and collected it 24 hours later. Fieldworkers hung 4.5 feet of sticky fly tape at least four feet from the ground near the latrine and food preparation area in a location away from smoke or stoves and protected from rain; they counted and speciated flies on the tape 24 hours later. Laboratory technicians enumerated *E. coli* and total coliform in water samples and *E. coli* and fecal coliform in toy rinses using membrane filtration.

### Outcomes

We pre-specified outcome measurement on ClinicalTrials.gov (#NCT02396407). We chose STH prevalence measured approximately 32 months post-intervention as the primary outcome of the spillover study because we believed spillovers were likely to impact STH transmission and because this objectively measured outcome is not subject to information bias. Stool samples with any ova were classified as positive. For each helminth, we quantified eggs per gram by multiplying the sum of egg counts from each of the duplicated slides by 12. We classified infection intensity into categories defined by the WHO based on the number of eggs per gram of stool (moderate/high intensity: ≥5,000 eggs/gram for *Ascaris*, ≥1,000 eggs/gram for hookworm, and ≥2,000 eggs/gram for *Trichuris*).^23^

Secondary outcomes included caregiver-reported 7-day diarrhea and respiratory illness prevalence measured approximately 28 months post-intervention. We defined diarrhea as caregiver’s report in the prior 7 days of 3+ loose or watery stools in 24 hours or 1+ stools with blood in 24 hours. We defined respiratory illness as a caregiver’s report in the prior 7 days of persistent cough or difficulty breathing.

While health outcomes serve as distal spillover effects, we also measured proximal spillover effects on environmental contamination after 32 months of intervention and WSH indicators after 28 months of intervention. Environmental contamination measures included the prevalence of *E. coli* and total coliforms in drinking water, the prevalence of *E. coli* and fecal coliforms in sentinel toy rinses, and the presence and number of synanthropic flies near the latrine and food preparation areas. WSH indicators included self-reported water treatment the day before the interview, storage of drinking water, presence of a latrine with a functional water seal, no visible feces on the latrine slab or floor, presence of a dedicated handwashing location with soap, and no visible dirt on the index child’s hands or fingernails.

### Sample size

We expected spillover effects to be smaller than effects on intervention recipients, so we powered the study to detect a relative reduction of 2.5-6% in primary outcomes, which was less than the 25% relative reduction expected in the WASH Benefits trial. We assumed prevalence differences for diarrhea (change from 14.2% to 8.2%), *Ascaris* (4.2% to 1.7%) and *Trichuris* (11.2% to 7.2%) and intra-cluster correlation coefficients (ICCs) ranging from 0.023 to 0.153 based on observational studies in rural Bangladesh and India.^24^ Assuming 80% power and a type I error of 0.05, we calculated the required sample size for each outcome of interest, adjusting for the ICC. Given these assumptions, the spillover study planned to enroll 2,000 children in 180 clusters (90 per arm).

### Statistical analyses

Two investigators (JBC, AE) independently analyzed primary and secondary outcome datasets following a pre-specified analysis protocol, which describes our analyses in full (https://osf.io/j9xht/). Here, we provide an overview of our analysis.

Analysis was intention-to-treat. Since WASH Benefits eligibility depended on pregnancy timing, we expected trial participants and adjacent neighbors to be equivalent on average except for their proximity to the WSH intervention, allowing us to make inferences about spillover effects by relying only on the cluster randomization. Our primary analysis estimated unadjusted prevalence ratios and differences for binary outcomes^21^ and unadjusted fecal egg count reduction ratios (1-ratio of mean intensity in intervention vs. control arm neighbors) for fecal egg counts. Our secondary analysis adjusted for covariates with bivariate associations with each outcome (likelihood ratio test p-value<0.2).^25^ We excluded binary covariates with prevalence <5%. We estimated parameters using targeted maximum likelihood estimation (TMLE) with influence-curve based standard errors accounting for clustering.^21^ Analysts were masked to intervention assignment until results were replicated.

We assumed children were missing at random and conducted a complete-case analysis. For outcomes with loss to follow-up exceeding 20% of the planned sample, we used TMLE to conduct an inverse probability of censoring-weighted analysis, which re-weights measured outcomes to reconstruct the original study population as if no children had missing outcomes.^26^

We assessed effect modification by pre-specified covariates: Euclidian distance to the nearest WASH Benefits compound, number of steps to the nearest WASH Benefits compound, presence of natural physical boundaries (e.g., pond) between spillover compounds and the nearest WASH Benefits compound, and the density of WASH Benefits compounds within a given radius of each spillover compound. All statistical analyses were completed using R version 3.2.3.

### Human subjects protection

We received approval from the Institutional Review Boards at the University of California, Berkeley (2011-09-3652), the International Centre for Diarrheal Disease Research, Bangladesh (PR-11063), and Stanford University (25863). Participation of human subjects did not occur until after written informed consent was obtained.

## Results

The spillover study screened 6,329 compounds neighboring WASH Benefits compounds for eligibility (Figure 3). Field workers enrolled 900 children in 90 clusters (“control neighbors”) and 889 children in 90 WSH clusters (“intervention neighbors”). Overall, 75% (N=634 control neighbors and N=710 intervention neighbors) of enrollees provided a stool specimen.

**Figure 3:**
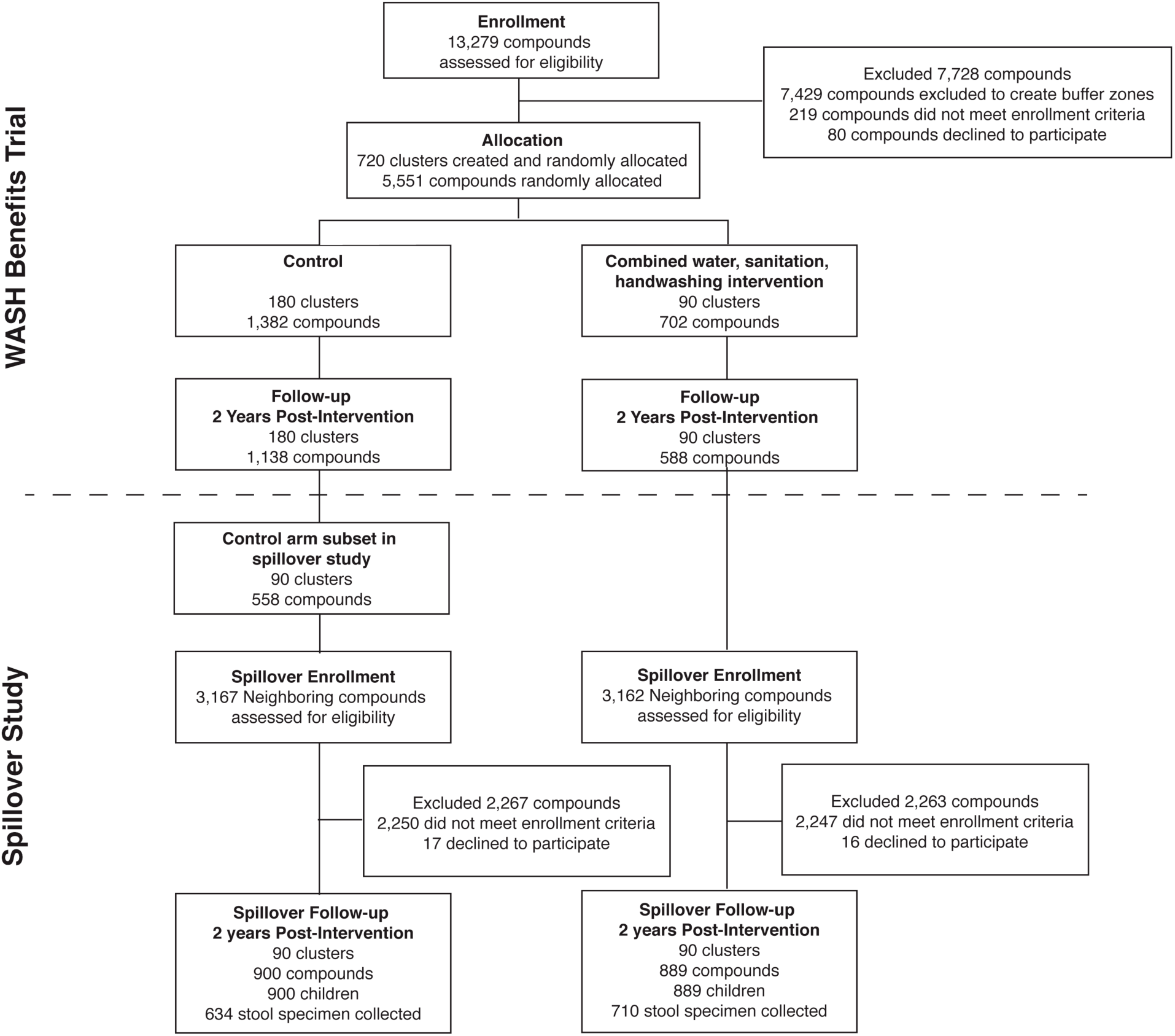
Participant flowchart

Characteristics of intervention neighbors and control neighbors enrolled in the spillover study were balanced by randomization, and neighbors’ characteristics were similar to those of WASH Benefits participants’ (Table 1). 815 (91%) intervention neighbors and 483 (54%) control neighbors knew of the WASH Benefits study (Appendix Table 1-2). Among intervention neighbors, 26% had spoken with WASH Benefits participants and 9% had spoken with WASH Benefits promoters about the study. While intervention adherence was high among WASH Benefits study participants at follow-up (Figure 4), there was no evidence of intervention use among intervention neighbors, control neighbors, and WASH Benefits control compounds at 2- year follow-up (Figure 4).

**Figure 4:**
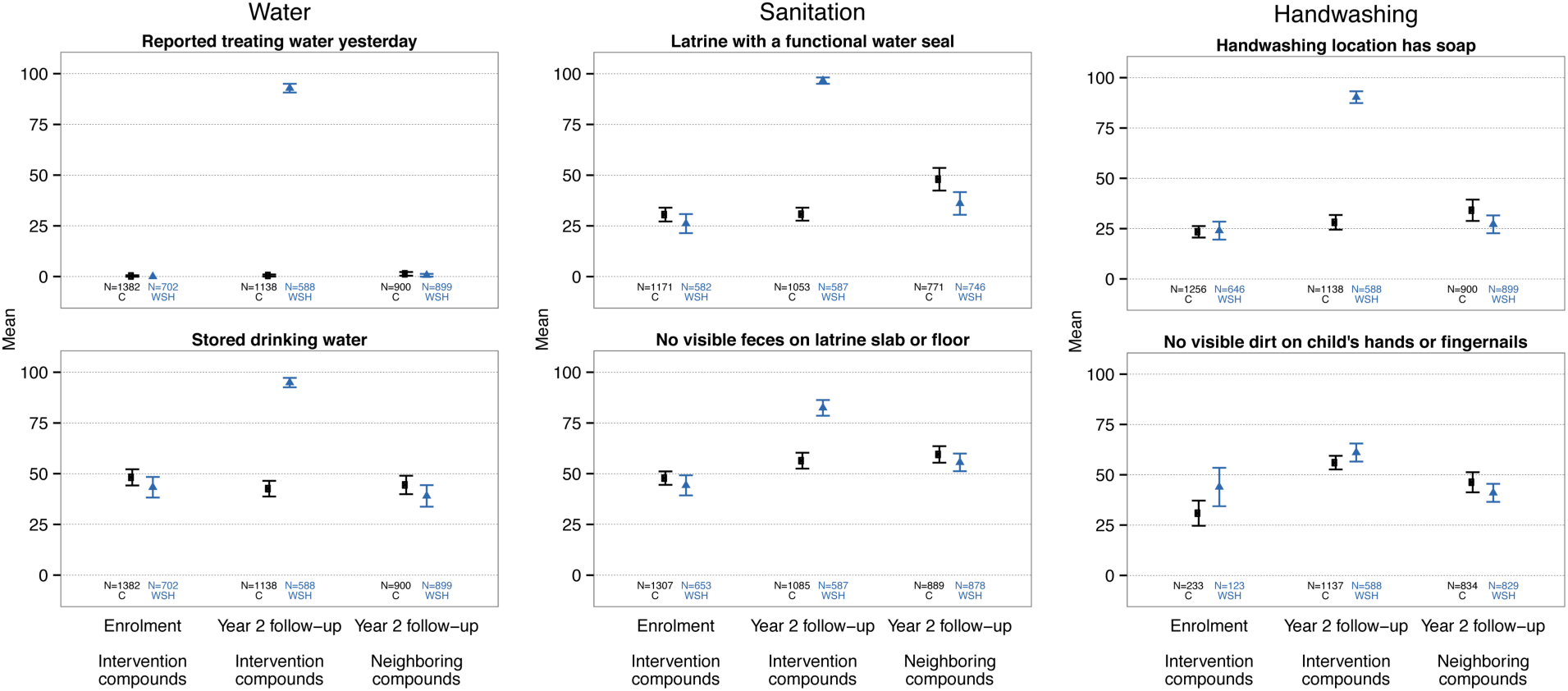
WASH indicators among WASH Benefits compounds and neighboring compounds.

**Table 1.**
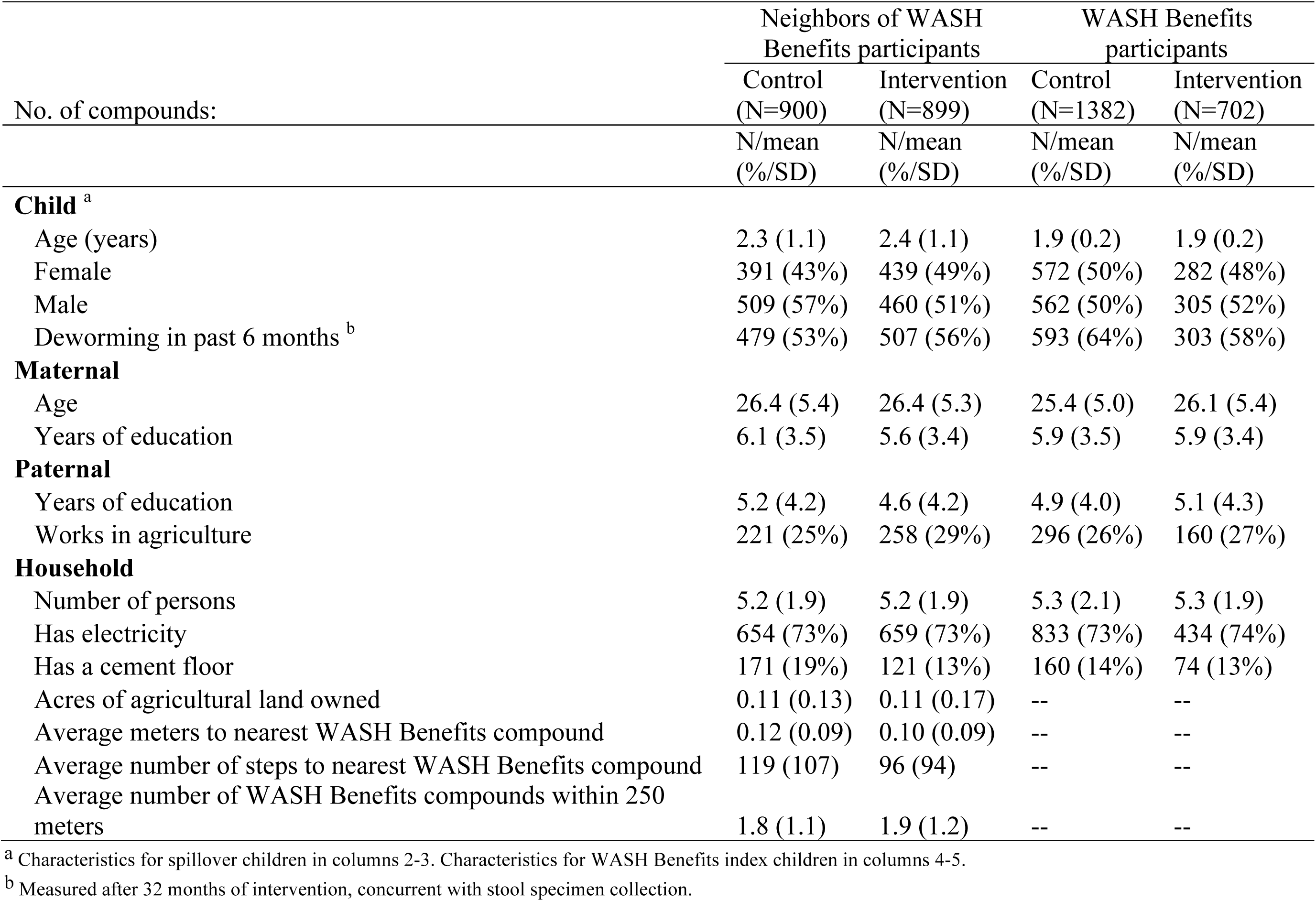
Spillover cohort enrollment and WASH Benefits trial participant characteristics by intervention group after 28 months of intervention

Median fly counts and *E. coli* and fecal coliform prevalence and mean log_10_ concentrations in sentinel toy rinses were similar between intervention and control neighbors (Table 2). The prevalence of *E. coli* detected in drinking water was lower for intervention vs. control neighbors (unadjusted prevalence ratio (PR)=0.88; 95% CI 0.80, 0.96). This effect was stronger among water samples collected directly from the tubewell (PR=0.83; 95% CI 0.73, 0.95), and there was no effect among samples from stored drinking water (PR=1.02; 95% CI 0.95, 1.10). The prevalence of total coliforms was similar between arms in all drinking water samples regardless of water source.

**Table 2.** Measures of environmental contamination among neighbors 32 months after intervention by intervention arm

|  | Control Neighbors |  |  | Intervention Neighbors |  |  | Unadjusted ratio of fly counts (95% CI) <sup>a</sup> |
| --- | --- | --- | --- | --- | --- | --- | --- |
|  | N Compounds | Median (SD) fly count | n (%) with any flies | N Compounds | Median (SD) fly count | n (%) with any flies |  |
| **Synanthropic flies <sup>b</sup>** |  |  |  |  |  |  |  |
| Near latrine | 717 | 3 (13) | 553 (78%) | 713 | 3 (21) | 576 (82%) | 1.16 (0.81, 1.66) |
| Near cooking area | 718 | 3 (23) | 570 (79%) | 711 | 3 (21) | 559 (79%) | 0.88 (0.64, 1.21) |
|  | N Compounds | Mean log<sub>10</sub> CFU/ 100 ml<sup>c</sup> (SD) | n (%) positive samples | N Compounds | Mean log<sub>10</sub> CFU/ 100 ml<sup>c</sup> (SD) | n (%) positive samples | Unadjusted prevalence ratio (95% CI) <sup>a</sup> |
| **Sentinel toys** |  |  |  |  |  |  |  |
| E. coli | 700 | 1.5 (1.3) | 558 (80%) | 697 | 1.5 (1.3) | 581 (83%) | 1.05 (0.99, 1.11) |
| Fecal coliforms | 700 | 3.4 (1.1) | 695 (99%) | 697 | 3.2 (1.2) | 691 (99%) | 1.00 (0.99, 1.01) |
| E. coli |  |  |  |  |  |  |  |
| All samples <sup>d</sup> | 718 | 0.9 (0.9) | 553 (77%) | 713 | 0.7 (1.0) | 481 (67%) | 0.88 (0.80, 0.96) |
| Samples from tubewell | 424 | 0.6 (0.9) | 281 (66%) | 470 | 0.4 (0.8) | 259 (55%) | 0.83 (0.73, 0.95) |
| Samples from stored water | 258 | 1.3 (0.8) | 238 (92%) | 219 | 1.5 (0.8) | 206 (94%) | 1.02 (0.95, 1.10) |
| Total coliforms |  |  |  |  |  |  |  |
| All samples <sup>d</sup> | 718 | 2.1 (0.5) | 710 (99%) | 713 | 2.0 (0.6) | 700 (98%) | 0.99 (0.98, 1.01) |
| Samples from tubewell | 424 | 1.9 (0.6) | 416 (98%) | 470 | 1.8 (0.7) | 457 (97%) | 0.99 (0.97, 1.01) |
| Samples from stored water | 258 | 2.3 (0.2) | 258 (100%) | 219 | 2.3 (0.1) | 219 (100%) | -- <sup>e</sup> |

<sup>a</sup> Standard errors account for clustering at the study cluster level.
<sup>b</sup> Includes *Musca domestica*, bottle flies (*Calliphoridae*), flesh fly (*Sarcophagidae*), lesser house fly (*Fannia canicularis*)
<sup>c</sup> For values below the detection limit (1 CFU per 100 mL for water, 2.5 CFU per 100 mL for toy rinses), we imputed 0.5 prior to taking the logarithm.
<sup>d</sup> Includes n=55 compounds who drew drinking water samples directly from a piped water source, which were not included a separate stratification category due to the low number of observations. 903 (63%) drinking water samples provided by participants were collected from tubewells, 487 (33%) were from stored water, 55 (4%) were from piped water, and 3 (<1%) were from water filters.
<sup>e</sup> Prevalence ratio could not be estimated because all samples contained total coliforms.

In the control arm, the prevalence of *Ascaris* was 31.4%, hookworm was 3.9%, and *Trichuris* was 3.9% (Table 3). There were no differences in STH prevalence between arms: *Ascaris* prevalence difference (PD)=0.00 (95% CI -0.07, 0.08); hookworm PD=0.01 (95% CI - 0.01, 0.04); *Trichuris* PD=0.02 (95% CI -0.02, 0.05); and any STH infection PD=0.02 (95% CI - 0.05, 0.09) (Table 3, Figure 5). There were no reductions in geometric fecal egg counts. Prevalence of moderate or heavy infections was <5% for all helminths among both intervention and control neighbors (Appendix Table 3). 4% of control neighbors (N=634) and 5% of intervention neighbors (N=711) were infected with more than one helminth. The prevalence was also similar between intervention vs. control arms for diarrhea (8.0% vs. 7.6%) and respiratory illness (8.6% vs. 9.2%) (Table 3).

Adjusted and inverse probability of censoring weighted analyses produced similar results for all outcomes (Appendix Table 4). For all outcomes, prevalence ratios and differences comparing neighbors of intervention vs. control were similar across levels of effect modifiers (Appendix Figures 1-7).

**Figure 5:**
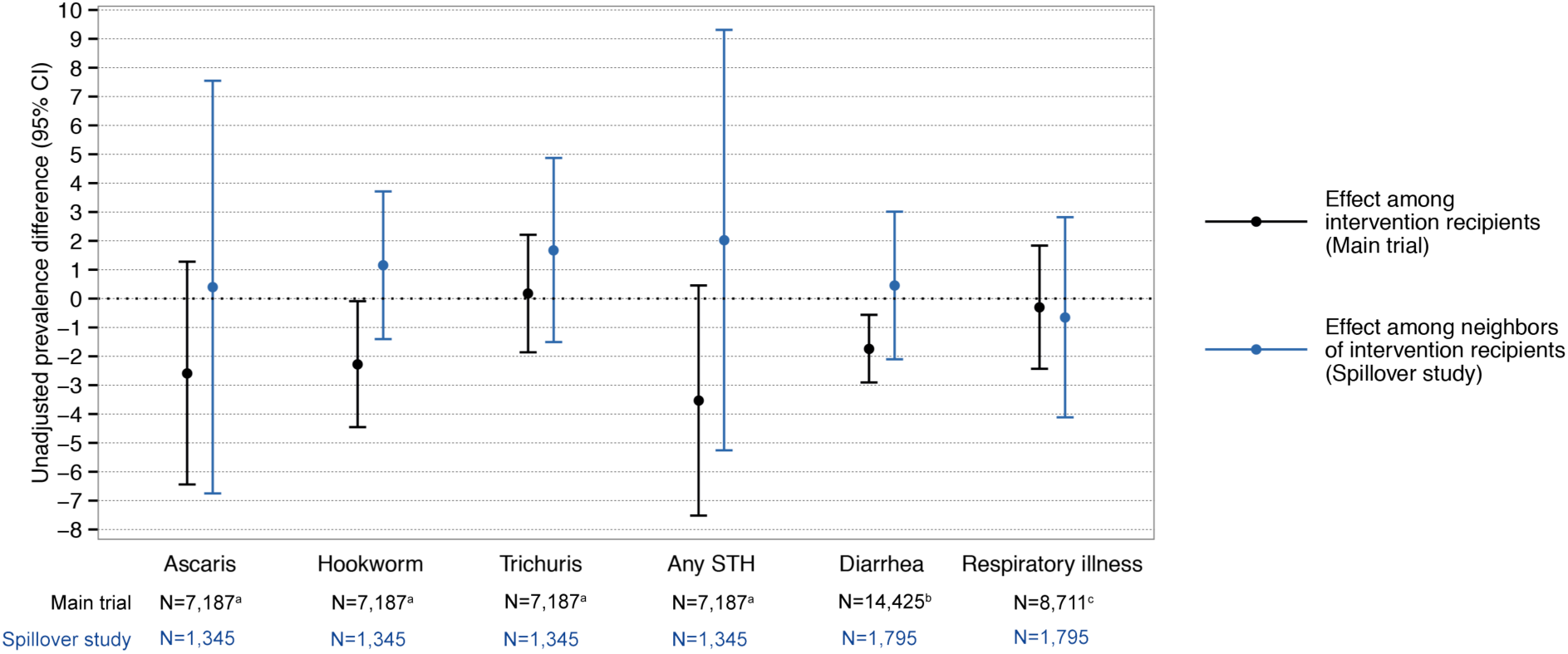
Comparison of prevalence differences for intervention vs. control among intervention recipients and their neighbors. ^a^Includes index children, pre-school age children, and school-aged children ^b^Includes children <36 months in the compound at enrollment ^c^ Includes index children and all other children under 5 years in the compound two years post-intervention

**Table 3.** Prevalence and unadjusted prevalence ratios and differences for diarrhea, respiratory illness, and soil-transmitted helminth infection among children neighboring WASH Benefits compounds after 32 months of WASH

|  | Control<br>Neighbors |  | Intervention<br>Neighbors |  | Unadjusted<br>prevalence ratio<sup>a</sup><br>(95% CI)<sup>b</sup> | Unadjusted prevalence<br>difference<sup>a</sup><br>(95% CI)<sup>b</sup> |
| --- | --- | --- | --- | --- | --- | --- |
|  | N | % | N | % |  |  |
| **Diarrhea prevalence** | 898 | 7.6 | 897 | 8 | 1.06 (0.76, 1.47) | 0.00 (-0.02, 0.03) |
| **Respiratory illness prevalence** | 898 | 9.2 | 897 | 8.6 | 0.93 (0.63, 1.37) | -0.01 (-0.04, 0.03) |
| **Soil-transmitted helminth prevalence** |  |  |  |  |  |  |
| *Ascaris lumbricoides* | 634 | 31.4 | 711 | 31.8 | 1.01 (0.81, 1.27) | 0.00 (-0.07, 0.08) |
| Hookworm | 634 | 3.6 | 711 | 4.8 | 1.32 (0.72, 2.42) | 0.01 (-0.01, 0.04) |
| *Trichuris trichiura* | 634 | 3.9 | 711 | 5.6 | 1.43 (0.75, 2.72) | 0.02 (-0.02, 0.05) |
| Any soil-transmitted helminth | 634 | 34.5 | 711 | 36.6 | 1.06 (0.86, 1.30) | 0.02 (-0.05, 0.09) |
|  | Control<br>Neighbors |  | Intervention<br>Neighbors |  | Unadjusted<br>prevalence ratio<sup>a</sup><br>(95% CI)<sup>b</sup> | Unadjusted prevalence<br>difference<sup>a</sup><br>(95% CI)<sup>b</sup> |
|  | N | Geometric<br>mean | N | Geometric<br>mean |  |  |
| **Soil-transmitted helminth infection intensity** |  |  |  |  |  |  |
| *Ascaris lumbricoides* | 634 | 3.23 | 711 | 3.92 | 0.16 (-0.27, 0.60) | 0.00 (-0.92, 0.93) |
| Hookworm | 634 | 0.21 | 711 | 0.24 | 0.02 (-0.11, 0.16) | -0.48 (-1.05, 0.10) |
| *Trichuris trichiura* | 634 | 0.2 | 711 | 0.32 | 0.10 (-0.09, 0.29) | 2.44 (-2.34, 7.21) |

<sup>a</sup> Prevalence ratios and differences compare the prevalence among intervention neighbors to the prevalence in control neighbors.
<sup>b</sup> Standard errors account for clustering at the study cluster level.
<sup>c</sup> Fecal egg count reduction ratio: (1-RR) x 100%, where the RR is the ratio of mean eggs per gram in the intervention vs. control arm.

## Discussion

We measured spillover effects of a combined WSH intervention on environmental contamination, hygienic behavior, and infectious outcomes in young children. By enrolling neighbors of randomly allocated trial participants, our study design enabled us to estimate geographically local spillover effects while relying on the original trial’s randomization for inference. Neighbors of intervention recipients were less likely to have *E. coli* detected in their tubewell water. However, there was no evidence of reductions in other measures of environmental contamination or of STH infection, diarrhea, or respiratory illness among intervention neighbors compared control neighbors.

Our environmental assessment measured proximal spillover effects on environmental contamination. We found lower *E. coli* concentration in tubewells of intervention neighbors compared to control neighbors. Though we did not find reductions in total coliforms in tubewell water, this indicator includes bacteria not of fecal origin^27^ and is less sensitive to changes in fecal input into the environment than *E. coli*. Improvements in latrine infrastructure may have reduced leakage into the groundwater;^28^ past studies have found fecal indicator bacteria in groundwater up to 2 m from pit latrines and up to 24.5 m in sandy soil.^29^ We did not find reductions in environmental contamination as measured by fly density, sentinel toys, and stored water, which capture surface level contamination. Together, these findings suggest possible spillover effects through groundwater but not surface level environmental contamination. Secondary contamination through poor hand hygiene, for example, may have counteracted improvements to source water quality.

Spillover effects may also have occurred if neighbors adopted interventions, but we found no evidence of intervention or behavior adoption. Limited hardware availability and lack of resources to purchase hardware likely inhibited diffusion of interventions to neighbors. Dual pit latrines would have been costly for neighbors to construct themselves, and Aquatabs^®^ and the water storage container delivered by WASH Benefits were not sold locally. Spillover effects may have been more likely if neighbors discussed interventions with WASH Benefits participants or saw them in use; however, only 26% of neighbors reported discussing WASH Benefits with intervention recipients. Finally, the absence of behavior change among neighbors may reflect limited knowledge of or perceived harm of illness or low social desirability of the WASH Benefits interventions.^30^

There are several features of this study that limit the generalizability of our findings. First, the intervention was only delivered to approximately 10% of each cluster on average. Studies have found that WSH interventions delivered to entire populations (e.g., introduction of municipal piped water and sewerage) were associated with reduced enteric infection.^14^–^17^ It is possible that a higher level of intervention coverage must be reached for WSH interventions to yield spillover effects. This is true for vaccines, many of which confer benefits on non-recipients once immunization coverage reaches a herd immunity threshold (typically over 75%).^7^ Second, the original WASH Benefits study found that the combined WSH intervention led to small reductions in STH prevalence (PD=-3.5%; 95% CI -7.5, 0.5) and diarrhea (PD=-1.7%; 95% CI - 2.9, -0.6).^20^ The size of spillover effects may be correlated with the size of effects on intervention recipients;^5^ in this study, impacts on intervention recipients may have been too modest to reduce transmission to neighbors.

Our study is subject to several limitations. First, we measured diarrhea and respiratory illness through caregiver report. Poor recall may have led to misclassification; however, because neighbors did not receive interventions, any misclassification was likely to be non-differential by study arm, which would have biased results towards the null. Second, double-slide Kato-Katz has low sensitivity in low infection intensity settings such as Bangladesh, where large-scale school-based deworming programs have been offered since 2008.^31^ This may have limited our statistical power to detect spillover effects, which we would expect to be smaller than effects on intervention recipients. Finally, we did not define social networks. A small body of evidence suggests that enteric and respiratory pathogens can spread through social networks;^32^ while few studies have examined this for WSH interventions, spillovers through social networks are theoretically plausible.^18^

## Conclusion

A compound-level combined WSH intervention reduced contamination of neighbors’ tubewell water but did not lead to spillovers for other proximal measures of contamination in the domestic environment or for child health outcomes. For proximal spillover effects to translate to distal spillover effects, improvements in neighbors’ health behaviors may have been necessary. Alternatively, spillover effects may be more pronounced in populations with higher disease transmission or higher levels of WSH intervention coverage in the community.

## Funding

This work was supported by the National Institute for Child Health and Human Development [1R21HD076216].

